# Meta-awareness, mind-wandering, and the control of ‘default’ external and internal orientations of attention

**DOI:** 10.1101/2024.09.13.612916

**Authors:** Isaac R. Christian, Samuel A. Nastase, Lauren K. Kim, Michael S. A. Graziano

**Affiliations:** Department of Psychology, Princeton University, Princeton NJ, 08544; Princeton Neuroscience Institute, Princeton University, Princeton NJ, 08544

**Author notes:** Corresponding Author: Isaac R. Christian.

**Keywords:** Mind-wandering, Meta-awareness, Default Mode, External Attention, Internal Attention

## Abstract

The “default mode” of cognition refers to the tendency to simulate internal experiences, rather than attending to external events in the moment. But in some contexts, external focus can become captivating enough to act as the default mode. To explore the relationship between prepotent internal and external default modes, we measured brain activity in forty participants using fMRI. Naturalistic movie clips were viewed, each one four times in sequence. When subjects were asked to focus attention on the videos, more mind-wandering events (distractions from the externally-focused task) occurred as the videos became less interesting with each repetition, and also when less engaging videos were presented. When subjects were asked to focus internally on breathing, more mind-wandering events (distractions from the internally-focused task) occurred when videos were most interesting (on the first repetition) and when more engaging videos were presented. In the fMRI data, inter-subject correlation, within-subject correlation, and GLM analyses found similar fronto-parietal networks engaged in transitions between default-controlled states regardless of the internal-external distinction, indicating more overlap in internal-external processing than previously assumed. We suggest that whether the default state is internal or external, and whether the sources that disrupt it are internal or external, depend on context.

## 1 Introduction

We are captured by the objects of our attention. Such objects can be external (e.g. a cute cat) or internal (e.g. a fond memory). Attention research has almost exclusively focused on characterizing the former, the dynamics of external attention. However, attention is often, if not more frequently, directed away from the external environment towards internal thoughts, memories, or plans (Verschooren & Egner, 2023). This ‘internal bias’ of attention is a result of the stable and predictable environment humans commonly find themselves in. Because we often engage in habitual routines (e.g. the same commute to work each day), it is more valuable to think about previous experiences and to plan for upcoming scenarios than it is to pay attention to predictable external events. As a result, the landscape of human thought is automatically tuned towards internal thoughts and periodically the strength of this internal pull is so strong that we may miss important cues in the environment, such as a sign on the highway.

Decades of neuroimaging have found one network to be associated with this kind of internal thought, named the Default Mode Network (DMN; Andrews-Hanna et al., 2010; Raichle et al., 2001). The default mode – the simulation of past, future and hypothetical experiences – is the common orientation of attention in the absence of an explicit task goal. In contrast, this network classically deactivates, while others come online, when the focus of attention turns towards external events and stimuli (Fox et al., 2005). The network most often associated with this kind of externally directed, sensory attention is the Dorsal Attention Network (DAN). The DAN enables intentional deployment of attention to features in the visual field (Vossel et al., 2014), controls visual working memory (Jerde et al., 2012), and exhibits a trend opposite that of the DMN, deactivating for tasks requiring internal attention (Corbetta et al., 2008). Another network, the Fronto-Parietal Control Network (FPCN), dynamically calls upon networks specializing in internal or external processes depending on the task demands (Dixon et al., 2018; Maillet et al., 2019; Spreng et al., 2010). While it is now well known that these networks are not exclusively involved in internally or externally oriented cognition (Smallwood et al., 2021; Yeshurun et al., 2021), it is considered established in the literature that there is a distinction between external and internal attention (Chun et al., 2011; Dixon et al., 2014) and that there is a neural basis for a default, internally oriented mode of cognition (Buckner et al., 2008).

This view of a default internal orientation, however, hides some complexity. It could be argued that “default” applies equally well to internal and to external modes, depending on circumstances. The brain naturally moves to one state or the other depending on which environment is more interesting. For example, when external stimuli have more attentional pull, then external attention becomes the default, and it takes extra cognitive effort to pull attention internally. Contexts in which external content can be just as captivating as internal thought are increasingly prevalent in the modern world: recent evidence shows that 25% of adults and 45% of US teens engage with social media ‘almost constantly’ (Perrin, 2018) and often face challenges disengaging from these sources (Holmgren & Coyne, 2017). Many have ventured that such uncontrolled engagement warrants classification as a mental health disorder, as uncontrolled use has been shown to produce detrimental effects on personal, social, and professional life (Bányai et al., 2017; Huang, 2022) as well as on cognition (Firth et al., 2019). Thus, internal and external sources can provoke similar uncontrolled modes of attention.

Mind-wandering is usually studied in tasks where people try to attend externally but wander to internally oriented cognition (i.e., stimulus independent thought; Smallwood & Schooler, 2015). In contrast, the study of meditation measures the ability of individuals to maintain attention on a single internal task while resisting the attraction of free, unconstrained internal mentation. In the meditation literature, therefore, mind-wandering is usually considered a transition from on-task internal content to off-task internal content (A. Lutz et al., 2008). Neither perspective typically considers the case when attention is supposed to be maintained on an internal task (e.g., thinking about a homework problem) but is distracted by external events (e.g., a siren) and control is required to resist the external distraction. Thus, “mind-wandering” as a term could just as fruitfully be applied to internally or externally oriented cognition. In some cases, the mind involuntarily drifts from desired external to distracting internal cognition, as is traditionally studied in mind-wandering. But in other cases, people may try to attend internally and become distracted by external events. The mind-wandering processes may be quite similar in both cases, or involve evolution of attention states according to similar principles.

Controlling one’s own attention to prevent mind-wandering involves at least two cognitive functions: first, becoming aware of the current state of attention, and second, directing attention based on that assessment (Clark & Dumas, 2015). The first function is often referred to as meta-awareness, or the explicit knowledge of the contents of thought (Schooler, 2002). It has also sometimes been conceptualized as a model of one’s own attention (Graziano and Kastner, 2011). Meta-awareness is believed to reduce the frequency of mind-wandering (Schooler et al., 2011). Meta-awareness naturally occurs intermittently in dynamic thought, but can be primed to occur more frequently if the task requires it. Sometimes people are asked to continuously monitor their attentional state for lapses, a method often denoted as ‘self caught’ (Chu et al., 2023). Another method is to give people intermittent reminders to use meta-awareness to check their attentional state (Murray et al., 2020).

In the present study, we asked two questions. First, can we find behavioral evidence that people do indeed switch attention between internal and external targets, and that the default, or “prepotent” state of attention switches from external to internal depending on circumstance? Second, can we find neural evidence from functional magnetic resonance imaging (fMRI) data that corroborates shifts of attention from automatic to controlled states, as well as neural markers indicating both the ‘meta-aware’ moments of mind-wandering and transient shifts from internal to external attention?

We presented participants with 2-minute clips from movies. In one condition, the external attention condition, we asked participants to attend to the movie and to press the button when they noticed that their attention had wandered off the movie (i.e., to self-report mind-wandering events; Figure 1A). In a second condition, the internal attention condition, we asked them to ignore the movie, attend to their own breathing, and press the button when they noticed their attention wandering off of their breathing (Figure 1B). In both attention conditions, subjects watched four repetitions of the movie, back to back. We expected the movie to become harder to attend with each repetition. We also tested six different movies, three categorized as “high salience” and three as “low salience.” In this way we varied the salience and interest value of the external stimuli to study the effect on attention and mind-wandering.

**Figure 1.**
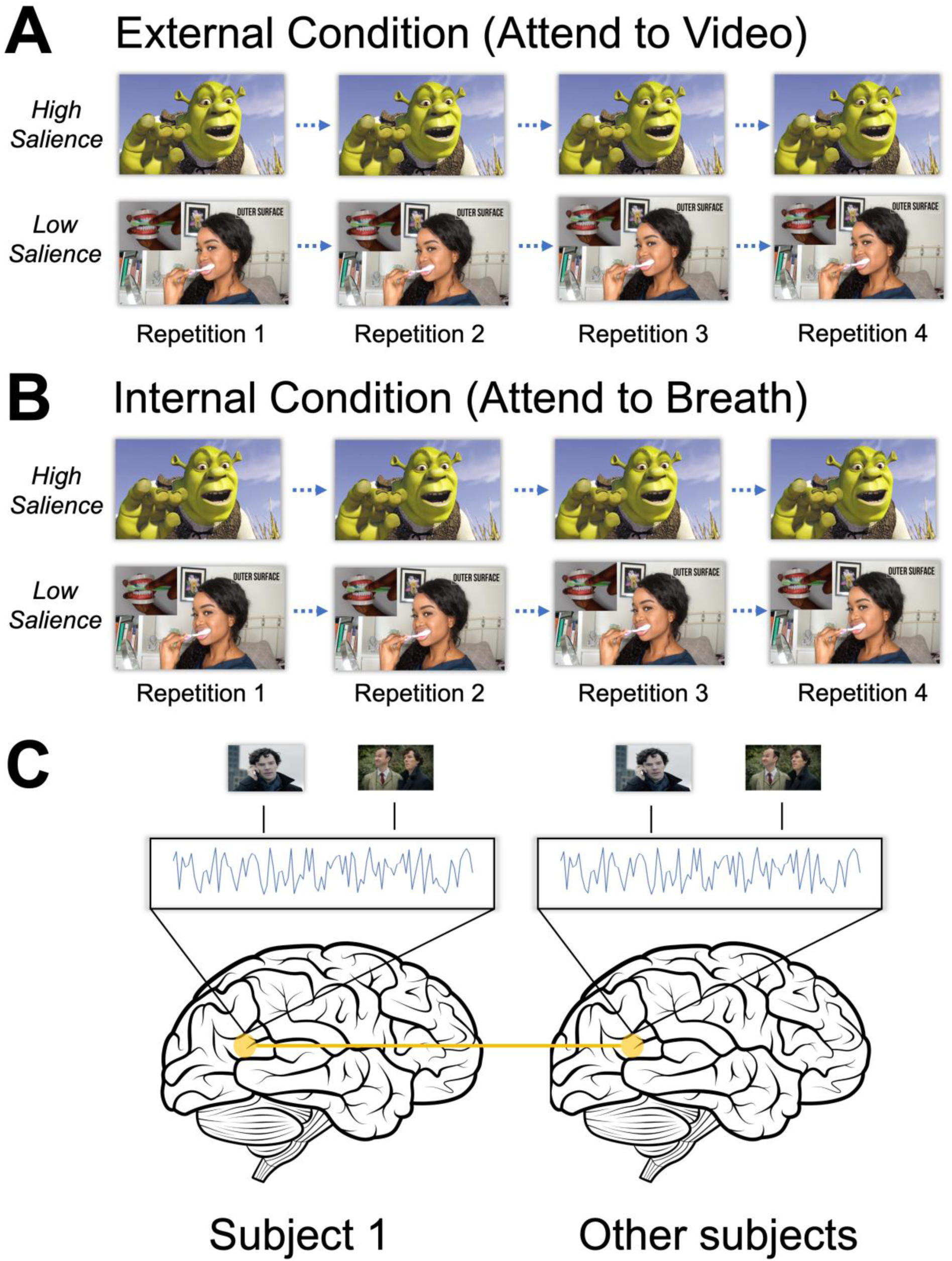
Experimental paradigm. Participants viewed a total of six videos during the experiment. Videos were categorized as high or low salience. Participants switched attention condition following three movies. **A.** In the external attention condition, a ∼2 minute video clip was presented on screen. Participants were instructed to pay attention to the video and press a button if they noticed that their mind had wandered from the video content. Participants repeated each task four times in a row, sequentially viewing the same video and indicating any lapses of attention. **B.** In the internal attention condition, participants viewed another ∼2 minute video clip, this time tasked with ignoring the external video and instead focusing on their breathing. Participants repeated each task four times in a row, sequentially viewing the same video and indicating any lapses of attention. **C.** Inter-subject correlation (ISC) was used to identify voxels that processed external information similarly across all subjects. To calculate ISC, the activation pattern of all voxels during stimulus presentation was extracted for one subject and correlated to the average timeseries across all other subjects for each respective voxel. This procedure was rotated for each subject. Subjects’ whole-brain ISC maps were averaged together to generate a final ISC image corresponding to voxels that exhibited similar time courses of activation across participants.

We hypothesized that in the external attention condition, maintaining attention on task would become more difficult with each repetition of the movie, and would also be more difficult for the low salience movies. This reduction of attention should be accompanied by more mind-wandering and thus a higher frequency of button presses.

In contrast, in the internal attention condition, when subjects needed to ignore the movie and focus on their breath, attention to the task should become easier with each repetition of the movie and should also be easier for low salience movies. This improvement in on-task focus of attention should be accompanied by a lower frequency of button presses. In this way, we should be able to track the dynamics of an external task with mind-wandering, and an internal task with external distractors inducing mind-wandering.

We further hypothesized that the effects of meta-awareness and control of attention would be measurable in cortical fMRI signals. The neural representation of the external video stimulus should be stronger and more widespread in the brain when the task is to pay attention externally, when the video is presented for the first time, and when the type of video is high salience. In contrast, the neural representation of the external video stimulus should be disrupted when the task is to pay attention internally, when the video has already been seen and is now being repeated, and when the type of video is low salience.

Moreover, by analyzing functional connectivity among brain areas and networks, it should be possible to track the change in how control and attention networks interact as the movie stimuli become less interesting over repeated presentations. We hypothesized that in the external attention condition, the movie would become harder to attend to over repetitions, and therefore the control and attention networks should show more interconnectivity, as the difficulty of control is increased. In contrast, we hypothesized that in the internal attention condition, the movie would become easier to ignore over repetitions, and therefore the control and attention networks should show less interconnectivity as the difficulty of control is decreased.

## 2 Methods

### 2.1 Subjects

Forty healthy human volunteers (aged 18-21, normal or corrected to normal vision, 37 right handed, 17 female) were recruited from the community and from a subject pool sponsored by Princeton University. All subjects provided consent and received course credit for participation or monetary compensation. One subject was excluded due to scanner malfunction. All procedures were approved by the Princeton Institutional Review Board.

### 2.2 Experimental Setup

Subjects lay supine on the MRI bed. Visual stimuli and video content were displayed using PsychoPy2 (Peirce et al., 2019) and projected onto a screen 80 cm from the eyes through an angled mirror mounted on top of the head coil using a digital light processing projector (Hyperion MRI Digital Projection System, Psychology Software Tools, Sharpsburg, PA, USA) with a resolution of 1920 x 1080 pixels at 60 Hz. Audio for each video was delivered with MRI-compatible Sensimetrics in-ear headphones. All responses were recorded using a button box held in the subject’s right hand and operated with the index finger.

### 2.3 Movie Stimuli

Six movie stimuli were used. Three originated from widely popular mainstream media sources, including a clip from BBC’s TV drama series Sherlock (2 m 19 s), Dreamwork’s animated comedy Shrek (2 m 7 s), and NBC’s sitcom the Office (2 m 25 s). In Shrek, the ∼2 minute clip consisted of a personal discussion between two characters about one character’s feeling of prejudice; in Sherlock, the scene comprised an initial synopsis of a crime scene; in the Office, three characters interacted comically in a skit. These movies were selected on the basis of their popularity and engaging story lines encompassing vulnerability, drama, and comedy, respectively. Our hope was that these videos would be highly engaging to participants, and for this reason we categorized them as ‘high salience’ videos. In contrast, the second set of three videos were instructional videos selected from YouTube: How to Make an Origami Heart (1 m 55 s), How to Bake a Cake (2 m 21 s), and How to Brush your Teeth (2 m 9 s). Our hope was that instructional videos on simple tasks would elicit less engagement compared to video content professionally designed to captivate audiences. We therefore categorized them as “low salience” videos. To measure the efficacy of high versus low salience categorization, at the end of the experiment participants were asked to rate how engaging each movie was on a scale from 1 to 10.

### 2.4 Task Design

The task design is shown in Figure 1. All subjects participated in two attention conditions. In the external attention condition, a video clip was presented on screen. Participants were instructed to pay attention to the video and press the button if they noticed that their mind had wandered from the video content. Mind-wandering was explicitly defined as content on mind that was unrelated to the content of the video, such as thinking about potential dinner plans or memories about the previous weekend.

Participants repeated this task four times in a row, sequentially viewing the same video and indicating any lapses of attention to the video with button presses. Once they finished four repetitions, participants were given an optional 30 s of rest and an opportunity to communicate with the experimenter. In the second condition, the internal attention condition, participants viewed another video clip (different from the video seen during the external attention condition), and this time were tasked with ignoring the video and instead focusing on the rhythmic sensation of their breathing, silently reciting “breathe in, breathe out” while keeping their eyes open and on screen. Once again, subjects were instructed to press the button if they noticed that their mind had wandered from their breathing, for example if their mind had wandered to the content of the video. The internal attention task was completed four times, with the same video presented each time. Thus one complete run of the external attention condition consisted of focusing on a video for four repetitions, and one complete run of the internal condition consisted of focusing on breath and ignoring an external video for four repetitions.

Participants completed three runs of the external attention condition and three runs of the internal attention condition. For each run, a unique video clip was used. All participants viewed the same six videos, but the attention condition (external, internal) and order of movie presentation were randomized for each subject. The six videos were pseudo-randomly sorted into two groups, with two high salience videos and one low salience video assigned to group A, and one high salience and two low salience videos assigned to group B. Half of participants completed the external condition using the three videos in group A and the internal condition using the three videos in group B. The other half of participants did the opposite, completing the internal condition using the three videos in group A and completing the external condition using the three videos in group B. Within each set of three videos, the order of videos was randomly varied between subjects. Finally, half of participants began the experiment performing three runs of the external attention condition followed by three runs of the internal condition, and the other half began with the internal attention condition followed by the external condition.

### 2.5 fMRI Data Acquisition

Functional imaging data were collected using a 3T MAGNETOM Skyra scanner (Siemens Healthineers AG, Erlangen, Germany), equipped with a 64-channel head/neck coil. Gradient-echo T2*-weighted echo-planar images (EPI) with blood oxygen level dependent (BOLD) contrast were used as an index of brain activity (Logothetis et al., 2001). Functional image volumes were composed of 40 near-axial slices with no interslice gap and an in-plane acceleration factor of 2 using Generalized Autocalibrating Partially Parallel Acquisition (GRAPPA), with slice thickness = 3.0 mm, field of view (FOV) = 200 mm, 80 x 80 matrix, 2.5 mm x 2.5 mm in-plane resolution, echo time (TE) = 30 ms, flip angle = 70°, and bandwidth of 1690 Hz/Px. A full functional volume was collected every 1.5 seconds (TR = 1500 ms).

Before functional runs, matching spin-echo EPI pairs with reversed phase-encode blips produced pairs of images with distortions going in opposite directions for blip-up/blip-down susceptibility distortion correction. An additional high-resolution structural image was collected for each participant, with 3D magnetization-prepared rapid acquisition gradient echo (MPRAGE) sequence, voxel size = 1 mm isotropic, FOV = 256 mm, matrix size = 256 x 256, 176 slices, TR = 2300 ms, TE = 2.96 ms, inversion time (TI) = 1000 ms, flip angle = 9°, iPAT GRAPPA = 2, bandwidth = 240 Hz/Px, anterior-posterior phase encoding, and ascending acquisition.

### 2.6 fMRI Preprocessing

Data were preprocessed using FMRIPREP version 1.2.3 (Esteban et al., 2019). T1-weighted volumes were corrected for intensity nonuniformity using N4BiasFieldCorrection v2.1.0 (Tustison et al., 2010) and skull-stripped using the OASIS template in antsBrainExtraction.sh v2.1.0. Spatial normalization through nonlinear registration to the ICBM 152 Nonlinear Asymmetrical template version 2009c (https://nist.mni.mcgill.ca/icbm-152-nonlinear-atlases-2009/) was completed using the antsRegistration tool of ANTs v2.1.0 (Avants et al., 2008). Brain tissue segmentation of cerebrospinal fluid (CSF), white-matter (WM) and gray-matter (GM) using Fast (Zhang et al., 2001) was performed on extracted T1w images.

Functional data were slice time corrected using 3dTshift from AFNI v16.2.07 (Cox, 1996) and motion corrected using mcflirt (FSL v5.0.9) (Jenkinson et al., 2002). Flirt (FSL) performed boundary-based registration with six degrees of freedom to co-register the corresponding T1w images to functional data (Greve & Fischl, 2009).

Motion correcting transformations, BOLD-to-T1w transformation, and T1w-to-template Montreal Neurological Institute (MNI) warp were concatenated and applied in a single step using antsApplyTransforms (ANTs v2.1.0) using Lanczos interpolation. All functional images were high-passed (0.007) using Nilearn’s signal cleaning function (https://nilearn.github.io/stable/modules/generated/nilearn.signal.clean.html). For further description of fMRIprep’s preprocessing pipeline see: https://fmriprep.readthedocs.io/en/latest/workflows.html. Before each movie repetition, a 7.5 second countdown was displayed on screen. This data was trimmed so that only TRs consisting of the movie presentation were used for analysis. Three buffer TRs were appended to the end of each repetition to account for hemodynamic lag.

### 2.7 fMRI Analysis: Inter-Subject Correlation

In conditions when participants attend to the external movie stimulus, the stimulus should influence their brain activity in a manner that is temporally correlated among the participants. In contrast, in conditions when participants are paying little or no attention to the movie stimulus, the stimulus should evoke less temporally correlated brain activity among subjects. Instead, cognitive processes unique to each subject will prevail and result in uncorrelated activity. As illustrated in Figure 1C, inter-subject correlation (ISC) was used to identify voxels that processed external information similarly across all subjects (Hasson et al., 2004; Nastase et al., 2019). To derive this measure, we used Brainiak’s Leave One Subject Out ISC function (Kumar et al., 2021). To calculate ISC, the activation pattern of all voxels during stimulus presentation was extracted for one subject and correlated to the average timeseries across all other subjects for each respective voxel. This procedure was rotated for each subject, iteratively calculating the correlation of one subject’s activation patterns to the group’s activation patterns. This analysis produced an ISC value for every held-out subject at every voxel.

We derived two complementary measures from these ISC maps: A group-level (thresholded) ISC map, which identified voxels whose activity was significantly correlated across subjects, and an unthresholded ISC measure, which provided an overall, descriptive estimate of how similar subjects’ neural responses were on average, without statistical testing. To identify voxels showing significant cross-subject correlations, we used a nonparametric bootstrapping method (https://brainiak.org/docs/brainiak.html). In each bootstrap iteration, ISC values from all held-out subjects were resampled with replacement and averaged, generating a distribution around the true mean ISC value for each voxel. After 10,000 iterations, voxel-wise hypothesis tests were conducted to identify ISC values significantly greater than zero. The resulting p-values were corrected for multiple comparisons using whole-brain FDR correction (p < .05). This analysis was performed for all repetitions in both the internal and external attention conditions, and results were averaged by salience type. To compute the unthresholded ISC measure, we averaged each subject’s held-out ISC map across all voxels to obtain a single value reflecting how strongly that participant’s brain activity aligned with the group overall. These subject-level ISC values were then averaged across participants, yielding a global measure of shared attention for each condition, repetition, which were again averaged by salience type.

### 2.8 fMRI Analysis: Within-subject connectivity Analysis

Within-subject functional connectivity is a way to measure correlations between the BOLD time courses of two different brain regions within the same individual. We used the Schaefer 200-parcel atlas (https://nilearn.github.io/dev/modules/description/schaefer_2018.html) to divide the cortex into regions. We computed pairwise correlations between all 200 regions, yielding a 200x200 matrix of Fisher-z transformed values. In principle, for each subject, a separate matrix could be computed for each for the 4 repetitions, for each of the 6 movies, resulting in 24 matrices.

To understand how connectivity may have changed as subjects experienced the same movies in repetition, we directly compared the first and last repetitions, in the following way. For each subject, we averaged together the three movies from the internal condition and the three movies from the external condition; then we averaged across subjects, averaging together all matrices for the internal attention condition and all matrices for the external attention condition. Finally, the aggregate matrices from the first repetition were subtracted from the aggregate matrices from the fourth repetition.

This subtraction yielded two group-level difference matrices. One showed how, in the internal attention condition, connectivity changed from the first to the last repetition; the other showed the same but for the external attention condition.

To assess statistical significance, we used a permutation-based approach. Prior to group-level averaging, we applied sign-flipping across subject-level matrices: a random vector of 1s and -1s (equal in length to the number of subject-movie matrices) was multiplied element-wise across matrices, flipping the sign of correlations for a subset of subjects. We then averaged these permuted matrices across subjects and movies to generate one null matrix per iteration. Repeating this process 10,000 times produced a null distribution of correlation matrices. We performed two-sample t-tests between each cell of the original group-level matrix and its corresponding distribution from the 10,000 permuted matrices. Resulting p-values were false-discovery-rate (FDR) corrected across all unique region pairs.

### 2.9 fMRI Analysis: GLM Analysis

To assess the activity associated with meta-awareness and the focus of attention, we constructed two event-related GLMs for the period prior to and after button presses. We hypothesized that the period immediately prior to the button press (0 to 3 s) would roughly capture episodes when meta-awareness was engaged and that the 0 to 3 s period following the button press would capture when participants redirected attention internally or externally following a mind-wandering event. We selected these 3s windows on the basis of previous work (Hasenkamp et al., 2012). To set up both GLMs, we concatenated each set of four repetitions into one functional run. This produced 6 functional runs per subject, one for each movie presented. The periods of interest (3 seconds before or after the button press) were convolved with a canonical hemodynamic response function (Glover, 1999) and inserted into a design matrix (one for each button press event) along with nuisance regressors. Nuisance regressors were defined as the first 5 principle components for CSF and WM masks as well as 3 translation and 3 rotation time series. Before fitting the model, functional time series were spatially smoothed using a 4mm FWHM gaussian kernel.

First level regression was performed using NiLearn’s FirstLevelModel function. The result was 12 beta maps for each subject, 6 for the period prior to the button press in each of the six movies, and 6 for the period following the button press in each of the six movies. We then averaged the beta maps of the three movies belonging to the internal attention condition and separately averaged the three movies belonging to the external condition. Thus, for each subject, there were two whole-brain beta maps for the meta-awareness period right before the button press (one for the external attention condition, one for the internal attention condition), and two beta maps for the period when the focus of attention shifted after the button press (again, one for the external and one for the internal attention condition).

A second-level model was fit using Nilearn’s SecondLevelModel, with the first-level contrasts of interest (external, internal, external minus internal) entered as input. To assess statistical significance while controlling for multiple comparisons, we applied a nonparametric permutation-based two-sided t-test (nilearn.glm.second_level.non_parametric_inference). This approach performs 10,000 random sign-flip permutations on whole-brain beta maps to generate a null distribution of voxelwise t-statistics under the null hypothesis of no effect. A voxelwise cluster-forming threshold of p < 0.001 was used. The resulting z-statistic map reflects the distribution of group-level effects corrected using permutation testing.

## 3 Results

### 3.1 Behavioral Results

Button presses indicated moments when participants believed their attention had wandered from the task at hand. More button presses suggested greater difficulty in attending to the target task; less button presses suggested less difficulty. Figure 2A shows the mean results averaged across all subjects, for all trials involving the high-salience movies (Sherlock, Shrek, and The Office). The graph shows the number of button presses, across four repetitions of the movie, for the external attention task and the internal attention task. A 2 X 4 analysis of variance (ANOVA) showed a significant main effect of external versus internal attention and a significant interaction (for external versus internal attention, F = 83.1, p < 0.001; for the interaction, F = 5.91, p < 0.001). A main effect indicating an overall change in button presses across the four repetitions was absent, due to the opposing patterns in which button presses decreased across the four repetitions for the internal attention condition and increased across the four repetitions for the external condition (F = 1.07, p > 0.05). To understand the specific pattern of results, more targeted statistical tests are described next.

**Figure 2.**
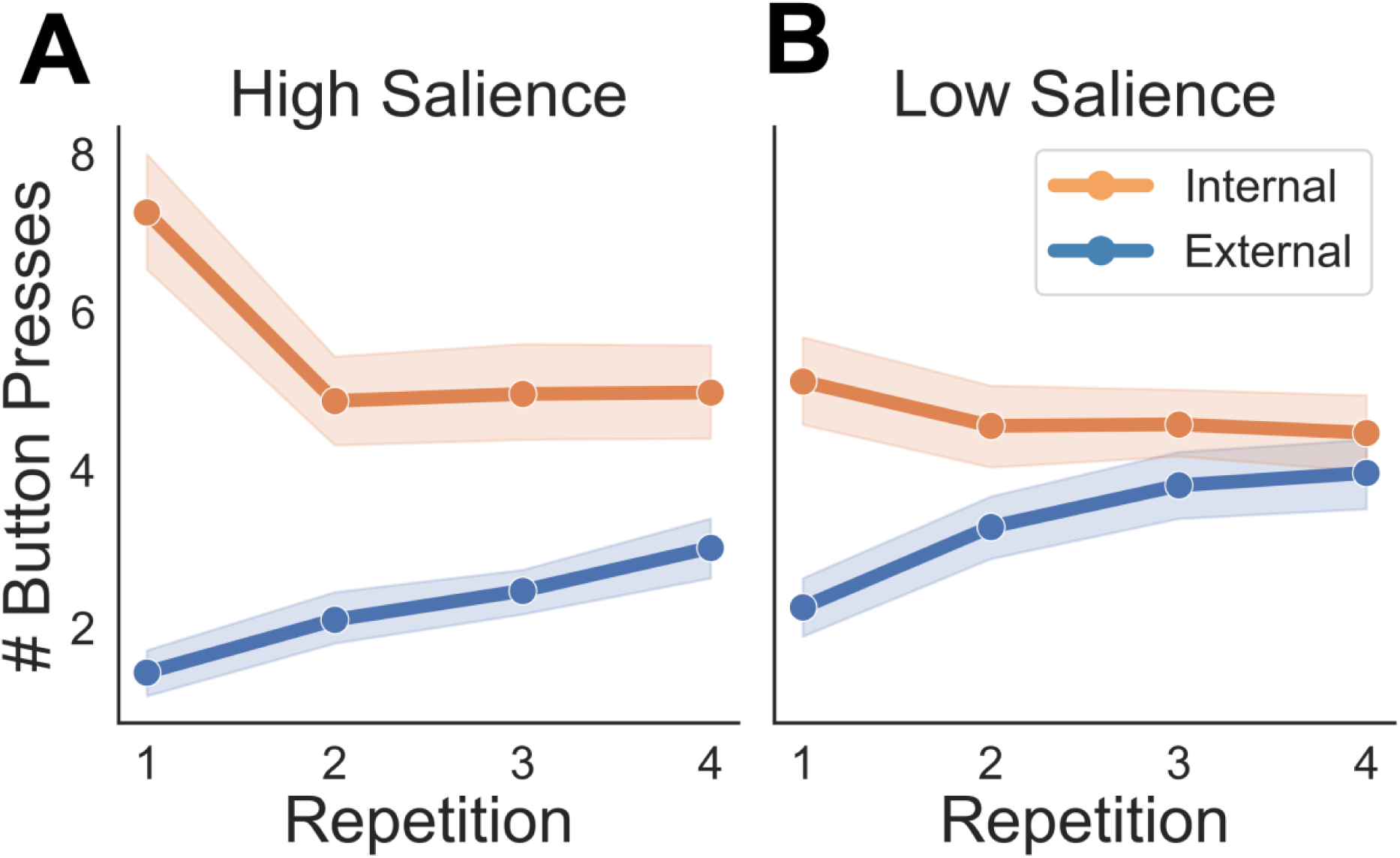
Number of self-reported mind-wandering events during attention tasks. **A.** Number of mind-wandering events for each repetition in internal (orange) and external (blue) attention conditions for movies categorized as high salience. Standard error is shown in lighter shade. **B.** Number of mind-wandering events for movies categorized as low salience.

In Figure 2A, in the external attention task (blue line), when subjects were asked to attend to the movie for the first time, they performed with few mind-wandering events (mean of 1.44 button presses), suggesting that they had little trouble keeping their focus on the movie. In each successive repeat of the movie, more mind-wandering events occurred, until, on the final repetition, subjects pressed an average of 3.02 times. This increasing trend was statistically significant (regression analysis on external attention data, t = 3.46, p < 0.001). As the movie became more familiar, the subjects appeared to find it harder to keep their focus on it and engaged in more mind-wandering.

In contrast, the orange line shows the results for the internal attention task, when subjects were asked to attend to their own breath. On the first viewing of the movie, subjects had difficulty keeping their attention away from the movie and maintaining attention on their breath, registering an average of 7.26 mind-wandering events. By the fourth presentation of the movie, subjects found it easier to focus on their own breath, pressing the button an average of 4.98 times. However, the change in attention was not gradual over the four repetitions. Instead, the shape of the curve suggests that on the first viewing, there was an initial draw to attend to the engaging movie, distracting subjects from their task and producing more mind-wandering events, whereas on subsequent repetitions of the same video, subjects were able to resist the distraction to a similar extent. Button presses during the first exposure to the movie were significantly greater than presses during the second, third, and fourth repetitions, but button presses were not significantly different among the second, third, and fourth repetition (t test between first and second repetition, t = 2.49, df = 112, p < 0.05; t test between first and third repetition, t = 2.35, df = 112, p < 0.05; t test between first and fourth repetition, t = 2.34, df = 112, p < 0.05).

The data in Figure 2A also suggest that the internal attention task was overall harder than the external attention task. Across all four repetitions of the movie, the number of button presses, and thus mind-wandering events, was higher in the internal task than in the external task. Even on the fourth presentation of the movie, when one might think the movie had become so boring that subjects would find it difficult to keep their attention on it (and therefore the external attention task should be harder), and easy to keep their attention on their own breath (and therefore the internal attention task should be easier), nonetheless, the internal condition still had significantly more button presses than the external condition (t = 2.97, df = 112, p < 0.01). This finding does not mean that internal attention is always harder than external attention. Instead, it suggests that in the context of this task, when subjects watched high salience movies, external attention on the movie was easier than internal attention on their own breath. In effect, in this context, external attention behaved more like a default mode.

Figure 2B shows the mean results for the low salience movies (instructional videos on origami, baking a cake, and brushing teeth). In some respects, the pattern is similar to that from the high salience movies, including a significant main effect in which button presses were overall greater for the internal versus the external attention condition (2 X 4 ANOVA, F = 17.53, p < 0.001), suggesting that the internal condition was again overall more difficult than the external condition; a significant interaction between repetition and attention condition (F = 2.64, p < 0.05); and no significant main effect of repetition (F = 0.59, p > 0.05). In the external attention condition (blue line), button presses were low on first presentation of the movie and increased with subsequent repetitions, with a significant increasing trend (regression analysis on external attention data, t = 3.04, df = 226, p < 0.01). In the internal condition (orange line), though button presses were higher on first presentation and lower on subsequent repetitions, the trend was not significant (t = -0.87, df = 226, p > .05). Subjects seemed relatively uniformly able to resist the distraction of the movie through all repetitions; we did not observe a difference in button presses between the first and subsequent repetitions (t test between first and second repetition, t = 0.75, df = 112, p > 0.05; t test between first and third repetition, t = 0.76, df = 112, p > 0.05; t test between first and fourth repetition, t = 0.90, df = 112, p > 0.05).

In a direct comparison between the high salience movies (Figure 2A) and the low salience movies (Figure 2B), some differences were observed. High salience movies appeared to be overall easier for subjects to attend to than low salience movies. For example, in the external attention condition, when subjects were asked to attend to the movie, they registered fewer mind-wandering events for the high salience movies than for the low salience movies (t test between high and low salience, in the external attention condition, t = 3.96, df = 454, p < 0.001). In the internal attention condition, the high salience movies were more distracting, making it harder for subjects to attend to their own breathing, as shown by the greater number of mind-wandering events for the high salience movies than for the low salience movies (t test between high and low salience, in the internal attention condition, t = 2.06, df = 454, p < 0.05).

Following the experiment, we asked participants to rate their level of engagement with each movie on a scale of 1 to 10. In total, we obtained 114 ratings for low salience movies (data available for 38 subjects X 3 movies) and 114 ratings for high salience movies. We found that self-reported levels of engagement were higher for the high salience videos (mean rating = 7.76) as compared the low salience videos (mean rating = 4.05). The difference was significant (df = 226, t = 13.14, p < 0.001). This result validates our separation of video stimuli into the high salience and low salience categories.

### 3.2 Neural Results: ISC

ISC is commonly used to identify representations of external stimuli that are shared across subjects, often described as stimulus evoked synchrony. We used ISC as a neural signature of attention to the external stimulus that was shared across subjects (Regev et al., 2019). In the external attention condition, when participants were explicitly tasked with paying attention to the movie, we expected shared attention to persist across repetitions of the movie, corresponding to higher ISC. In the internal condition, we expected that ignoring the external stimulus would produce less shared representation of the external stimulus across all four repetitions, corresponding to lower ISC. Additionally, we expected that high salience movies would be easier to focus on in the external attention condition, and harder to ignore in the internal attention condition, as compared to the low salience stimuli, and that these differences would also be reflected in the ISC data.

Figure 3A and B show an initial analysis of the ISC results, in which unthresholded ISC values (average ISC value across all voxels, across all subjects, across high or low salience movies) are displayed. Figure 3A shows data for the high-salience videos. Here, the effect of repetition is clear. For both the external attention condition (blue bars) and the internal attention condition (orange bars), the first presentation of the movie, relative to the other repetitions, corresponded to the greatest ISC, indicating the greatest amount of inter-subject correlation and therefore the greatest amount of cortical processing of the movie. Each subsequent repetition was associated with decreasing ISC, indicating a decreasing processing of the stimulus details, as the subjects’ attention on the movie waned and more mind-wandering events occurred. In addition, the external attention condition (blue bars) showed consistently higher ISC values than the internal attention condition (orange bars). This difference suggests that subjects were successful at reducing attention to, and processing of, the movie stimulus in the internal attention condition, when they were tasked with attending to their own breath. Figure 3B shows data for the low-salience videos. Here similar trends were observed.

**Figure 3.**
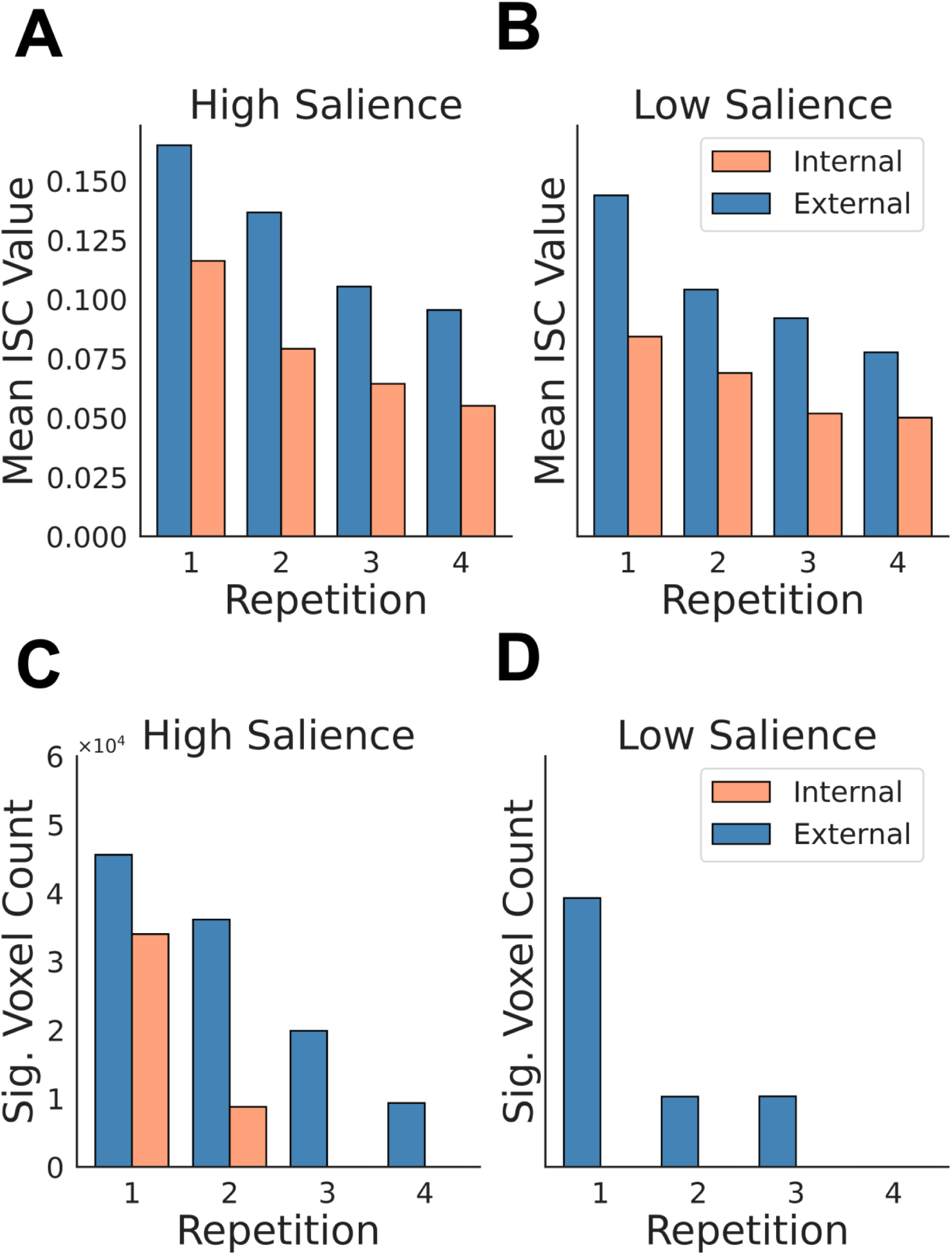
Unthresholded and thresholded inter-subject correlation. The top row shows unthresholded mean ISC values across subjects, where each bar represents the average of all subjects’ whole-brain ISC maps prior to statistical testing. The bottom row shows the group-level statistically significant results, where each bar represents the total number of statistically significant voxels identified using a nonparametric bootstrapping procedure. Blue bars indicate repetitions when attention was directed externally on the video; orange bars indicate repetitions when attention was focused inwardly on the breath. Results for each of four repetitions are displayed. **A.** Unthresholded ISC during viewing of high-salience movies. **B.** Unthresholded ISC during viewing of low-salience movies. **C**. Number of significant ISC voxels during high-salience movie viewing. **D.** Number of significant ISC voxels during low-salience movie viewing.

A mixed linear effects model confirmed the two most obvious trends in the data: the unthresholded mean ISC decreased with each repetition irrespective of attention condition or salience (main effect: z = -17.69, p < 0.001) and differed between attention condition, irrespective of repetition or salience (main effect: z = -14.48, p < 0.001). Mean ISC also decreased at a quicker rate in the external condition than in the internal condition (two-way interaction between repetition and condition: z = 4.17, p < 0.001).

Although in comparing Figure 3A to 3B, it appears that the high-salience movies evoked slightly higher ISC values than did the low-salience movies, this difference was not statistically significant (z = -0.50, p > 0.05). At least when measured by the unthresholded, brain-wide ISC values, shared external attention was consistent regardless of the kind of movie presented. This lack of a statistical difference is contrary to our original hypothesis: we expected that mean ISC would mimic results from behavioral analysis, showing privileged representation of videos that were considered more interesting and harder to ignore.

However, the brain-wide mean of unthresholded ISC values provides only a rough measure of external attention. Some voxels, those with high ISC values, are presumably engaged in processing stimulus content, while other voxels, those with low ISC, are presumably less involved in processing the stimulus. Including all voxels together in a brain-wide average introduces noise and potentially masks signal if the goal is to accurately estimate sensory encoding. We therefore performed a second ISC analysis, shown in Figure 3C and 3D. Here we plot the number of voxels that, when tested among all subjects, showed a statistically significant ISC. Here the same trends are evident, but the effect of movie salience is more pronounced. We did not, however, perform statistics on this second method of plotting the ISC results, since bootstrapping methods did not lend themselves to a metric of the total number of significant ISC voxels.

### 3.3 Neural results: Connectivity Analysis

In the external condition, we expected that remaining on task would become more challenging as participants repeatedly watched a movie clip, requiring an increase in the coordination of attention control networks to remain focused externally. By hypothesis, cortical areas involved in attention control should show greater functional connectivity as the movie clip was repeated more times. To test this hypothesis, we calculated connectivity matrices by correlating the time course of each brain region to the time courses of all other regions, using a 200-region parcellation of the cortex (see Methods). We subtracted the correlation values of the first repetition (when keeping attention on the movies naturally required the least control effort) from the correlation values of the fourth repetition (when keeping attention on the movies naturally required the most control effort).

Figure 4A shows the results for the external attention task. A red color in the matrix indicates a case in which a functional connection between two brain areas became stronger from the first to the fourth repetitions of viewing movies. A blue color, in contrast, shows a case in which a functional connection between two brain areas became weaker after four repetitions. There were a total of 99 connections that increased significantly from run 1 to run 4 (see Methods for statistical significance testing).

**Figure 4.**
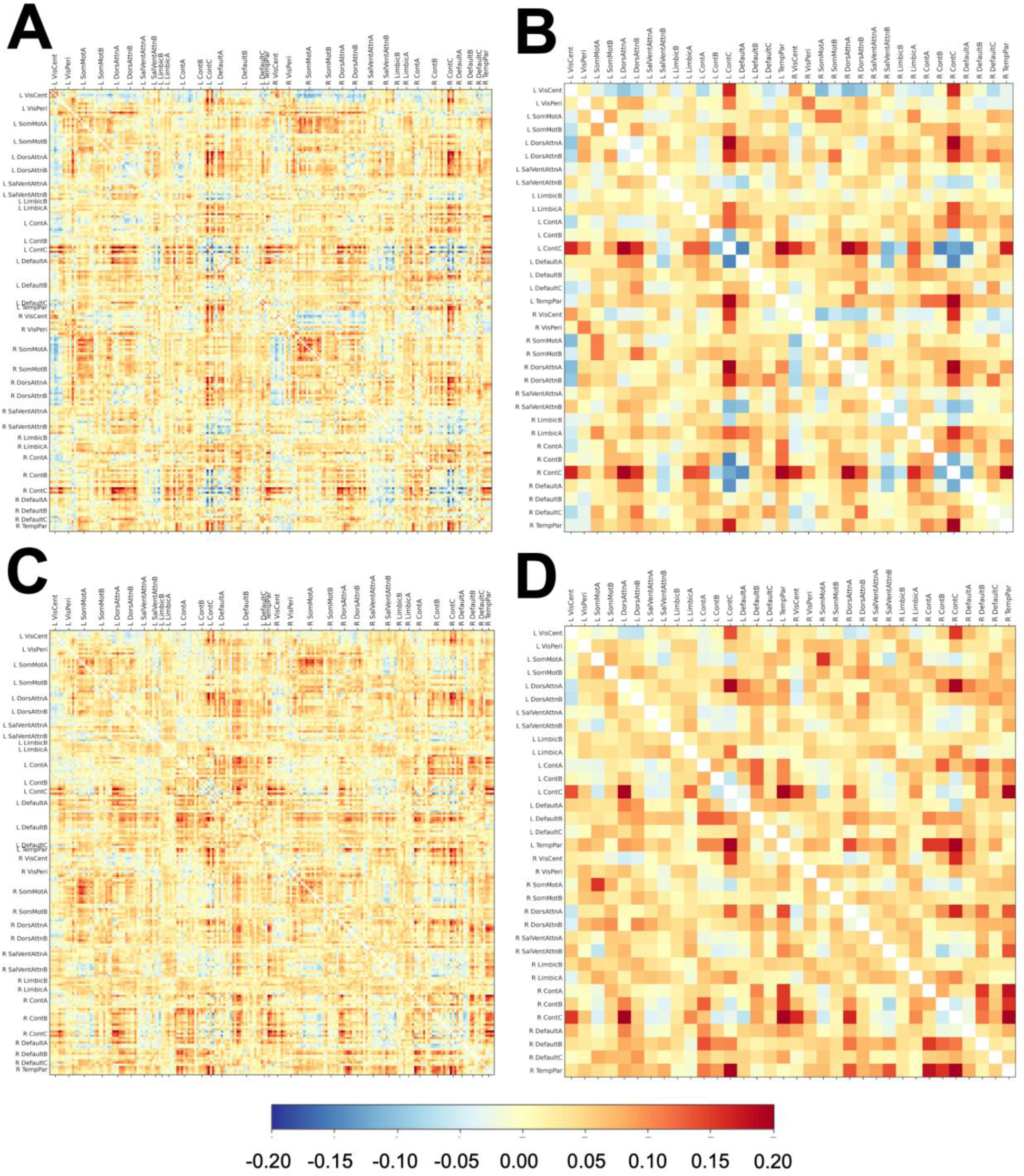
Functional connectivity analyses by region and network. Positive values (red) indicate correlation between pairs that increased from run 1 to run 4, and negative values (blue) indicate correlations that decreased. **A.** The connectivity of 200 regions (e.g., Region A correlated to Region B, C, etc..) in repetition 1 subtracted from the connectivity of 200 regions in repetition 4, for the external condition, averaged across movies. **B.** The connectivity of 34 networks in repetition 1 subtracted from the connectivity of 34 networks in repetition 4, for the external condition, averaged across movies. **C.** The same as A but for the internal condition. **D.** The same as B but for the internal condition.

Figure 4A shows so many connections between brain areas that the pattern can be difficult to interpret. One way to bring clarity is to consider cortical networks, rather than individual cortical areas. In the standard 200-area parcellation of the cortex used here (https://nilearn.github.io/dev/modules/description/schaefer_2018.html), the 200 areas have been assigned to 34 cortical networks, 17 in each hemisphere. The networks range from the smallest, containing 2 cortical areas, to the largest, containing 13 areas. Figure 4B shows connection pairs between networks, rather than between individual areas. To characterize the strength of the connection between two networks, we used the following method. If network A included N areas, and network B included M areas, then a total of NxM possible connections can exist between the networks. For each connection we measured a connection strength, then averaged the results across all MxN connections to arrive at a mean strength of connection between network A and B. We then subtracted the connection strength for repetition 1 from the connection strength for repetition 4. The result is plotted in Figure 4B.

Table 1 lists all 18 pairs of networks that, in the external condition, showed a significant change in connectivity from the first run of the movie to the last. In every case, the change was an increase. The interaction between these networks is presumably involved in controlling and maintaining attention on the movies as the movies become less interesting through repetition and therefore the task of attention control becomes more difficult. Among the many pairs of networks shown in this table, note, in particular, the increased connectivity between components of the DAN and the

**Table 1.**
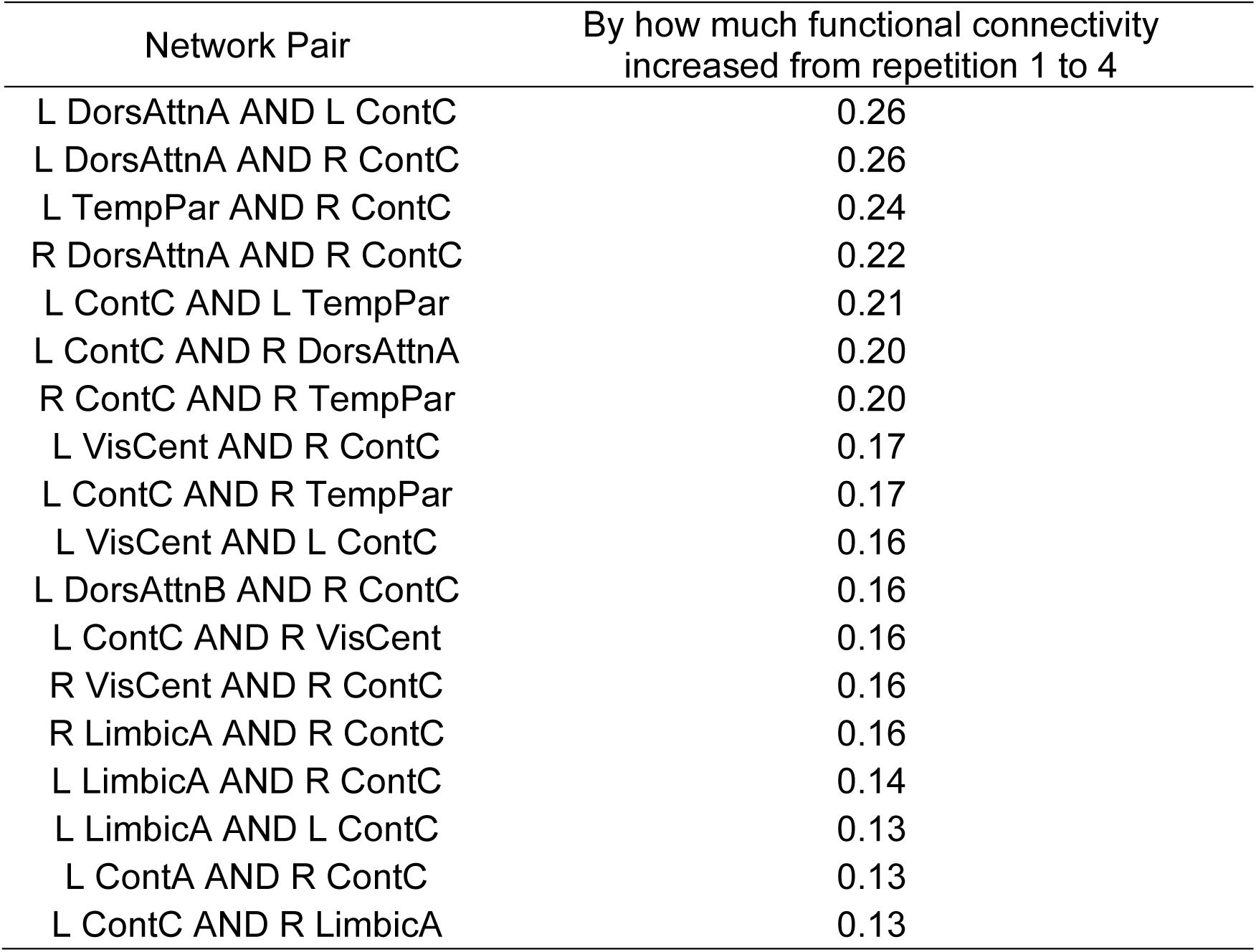
Connections showing significant increases in functional connectivity from repetition 1 to repetition 4 in the external condition. Functional connectivity was computed separately for repetitions 1 and 4. The difference (repetition 4 minus repetition 1) reflects increases in connectivity as the task became more challenging to attend. Significance was assessed against a permuted null distribution, and surviving connections are listed in the table. Column 1 lists the network pairs, and Column 2 reports the magnitude of the connectivity increase. Connections are sorted in descending order of connectivity change.

FPCN (e.g. between LDorsAttnA and LContC). We also found increased connectivity between components of the FPCN and networks previously associated with internal, rather than external, attention (e.g. between RContC and R TempPar). Increased connectivity between control networks and those typically associated with internal attention, therefore, may reflect a greater demand for internal attention processes. As the external environment becomes less exciting and the tendency to mind wander increases, monitoring one’s attention state becomes increasingly important. In that speculation, maintaining external attention is more complex than it may seem on the surface, requiring both external attention on the stimulus and internal attention on one’s own attention state, recruiting complex interactions between networks involved in control, external attention, and internal attention.

Figure 4C shows the results of a similar connectivity analysis, but applied to the internal attention condition. Here, subjects were asked to attend to an internal state, their own breathing, while monitoring their attention. The figure once again shows pairs of areas in the 200-area parcellation. For each pair, a connection strength was computed for the first and the last repetition of each movie; then the connection strength for the first run was subtracted from the connection strength of the fourth run, thereby indicating the amount that the connection changed over the course of the four repetitions. Connections that increased (red) outnumber connections that decreased (blue).

Figure 4D shows the result, for the internal attention condition, for connections among the 34 cortical networks, rather than among the individual 200 areas. Table 2 lists all 10 pairs of networks that, in the internal condition, showed a significant change in connectivity from the first repetition of the movie to the last. In every case, the change was an increase in connectivity. This overall increase in connectivity was surprising to us. In the internal attention condition, as the movie repeated and therefore became less interesting, we predicted that subjects would find it easier to focus internally on their own breathing. As noted above, the behavioral data supports that interpretation.

**Table 2.** Connections showing significant increases in functional connectivity from repetition 1 to repetition 4 in the internal condition. Functional connectivity was computed separately for repetitions 1 and 4. The difference (repetition 4 minus repetition 1) reflects increases in connectivity as the task became more challenging to attend. Significance was assessed against a permuted null distribution, and surviving connections are listed in the table. Column 1 lists the network pairs, and Column 2 reports the magnitude of the connectivity increase. Connections are sorted in descending order of connectivity change.

| Network Pair | By how much functional connectivity increased from repetition 1 to 4 |
| --- | --- |
| L TempPar AND R ContC | 0.24 |
| R ContC AND R TempPar | 0.23 |
| L DorsAttnA AND R ContC | 0.21 |
| L ContC AND L TempPar | 0.21 |
| L ContC AND R TempPar | 0.20 |
| R ContA AND R TempPar | 0.18 |
| R VisCent AND R ContC | 0.16 |
| L TempPar AND R ContB | 0.15 |
| L LimbicA AND L ContC | 0.12 |
| L LimbicA AND R ContC | 0.10 |

Therefore, we hypothesized that as the task became easier over the course of the four movie repetitions, the engagement of attention and control networks should become reduced. In contrast, the data show a clear increase in connectivity. In particular, resembling the case for the external attention condition, increased connectivity was found between components of the DAN and the FPCN (e.g. L DorsAttnA and R ContA), and between the control network and networks associated with internal attention (e.g. between R ContC and R TempPar). This pattern suggests that the task of controlling and maintaining attention on breath required more, rather than less, engagement of the crucial attention and control networks over time. It is possible that, although the movie distraction became weaker as the movies became less interesting, the task of maintaining focused internal attention may nonetheless have required a ramping up of control over time.

Figure 5 provides another way of comparing the external and internal conditions. Here, the color code indicates which statistically significantly connections between networks increased from the first to the last repetition, for both the external and internal attention conditions (green), just the internal attention condition (red), just the external attention condition (blue), and neither (black). The blue dots prevail. This result suggests that in the external condition in particular, when subjects were trying to maintain attention on a movie through all four repetitions, a set of connections in the cortex became stronger as the effort of maintaining attentional focus became harder; and whereas a similar trend occurred in the internal attention task, the effect was weaker, involving fewer connections.

**Figure 5.**
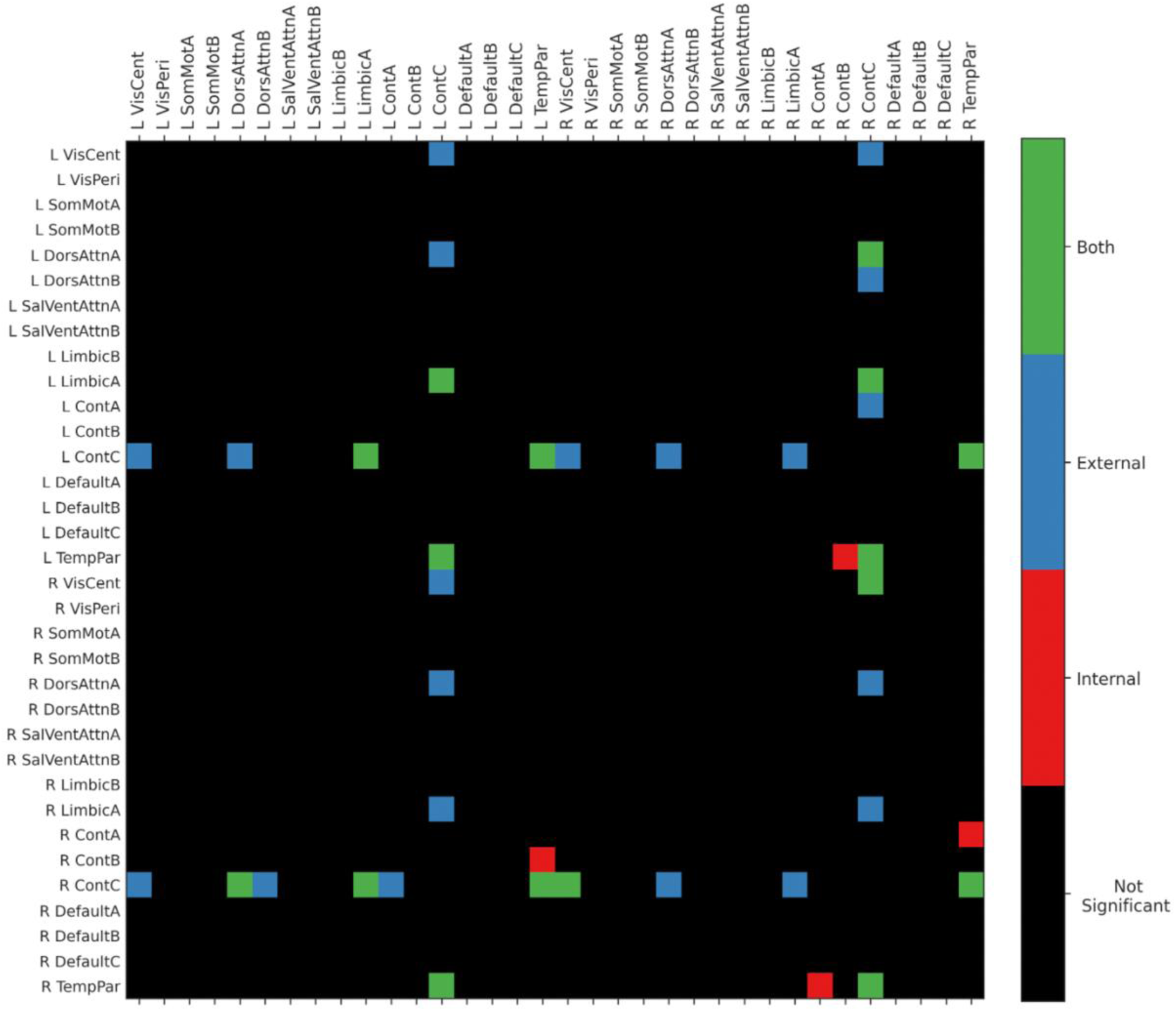
Network pairs showing significant increases in functional connectivity from repetition 1 to repetition 4 in External and Internal conditions. Red dots indicate connections that increased from run 1 to run 4 in the internal condition, blue dots show the same but for the internal condition, and green dots show connections that increased for both internal and external conditions. The remaining cells (black) did not significantly increase in internal or external conditions.

### 3.4 Neural Results: General Linear Model

We constructed four distinct general linear models (GLMs). The first two GLMs targeted brain activation during the 3 seconds preceding each button press: one GLM for the internal attention task, and one for the external attention task. This time window was intended to capture the period when participants introspected on their attentional state and evaluated whether it matched their intended focus, following the interpretation of previous studies (Christian et al., 2025; Hasenkamp et al., 2012). In that time period, participants are presumed to have formed a second-order, metacognitive representation of their own mental content, explicitly directing awareness toward their state of awareness at some point in this 3 s window.

We found that the networks engaged during meta-awareness were largely consistent across internal and external tasks (Figure 6A, 6B; see supplementary information for a table with cluster-level information). The most robustly engaged regions included the salience network, FPCN, and DMN. We also performed a subtraction (external attention condition minus internal attention condition). As shown in Figure 6C, almost no significant activity remained, indicating a near-complete overlap in the activation patterns for the two tasks. However, some condition-specific effects were observed. Orbitofrontal cortex and paracingulate cortex showed greater activation in the external attention task, whereas inferior temporal cortex, lateral occipital cortex and superior parietal lobule were more active in the internal attention task.

**Figure 6.**
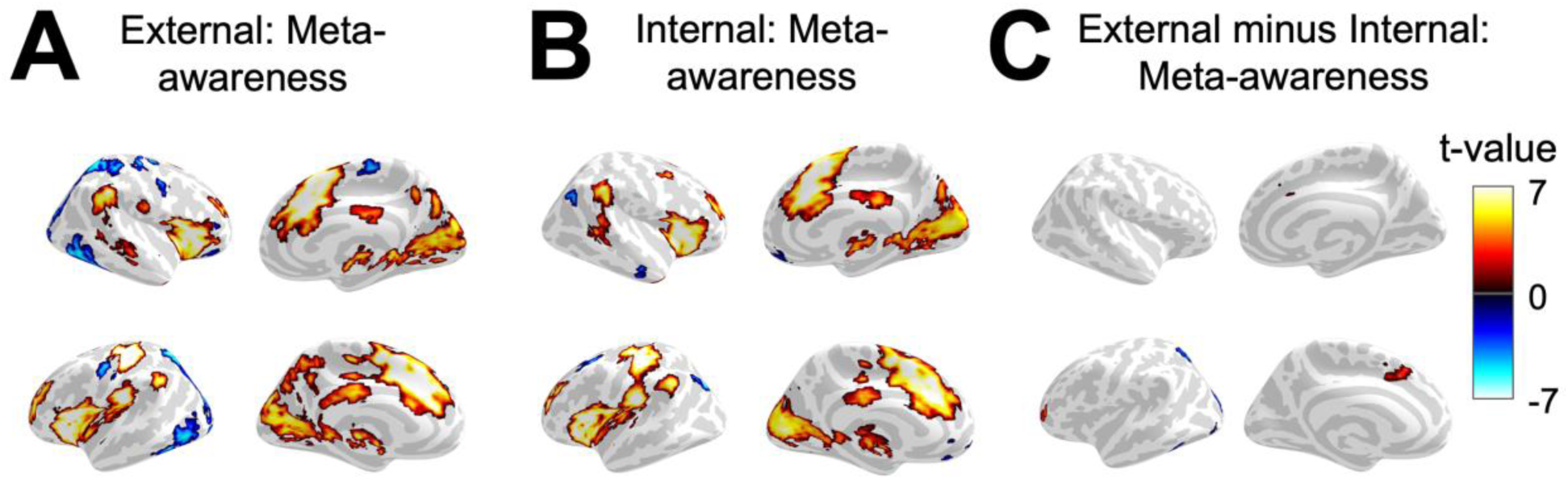
Voxel-wise univariate analysis for task periods involving meta-awareness. The group-level GLM estimated activation map (regression coefficients) for the 3 s period prior to the button press was extracted and projected onto the cortical surface. For panels A and B, positive t-values indicate voxels more strongly activated for the meta-awareness period and negative t-values indicate voxels that were less so; for panel C, positive values indicate voxels more active for the external compared to internal condition, while negative values indicate the opposite. **A.** The activation map for the meta-awareness period in the external condition. **B.** The activation map for the meta-awareness period in the internal condition. **C.** Subtracting the activation map for meta-awareness in the external condition from the period of meta-awareness in the internal condition.

The second set of GLMs examined networks involved in reorienting attention during the 3 s period immediately following button presses (one GLM for the internal attention task, and one for the external attention task). Once again, the networks engaged were similar across internal (Figure 7A) and external (Figure 7B) tasks (see supplementary information for a table with cluster-level information). When we performed a subtraction (external attention task minus internal attention task, Figure 7C), only one cluster remained, located in the posterior cingulate cortex, displaying stronger activation for internal attention.

**Figure 7.**
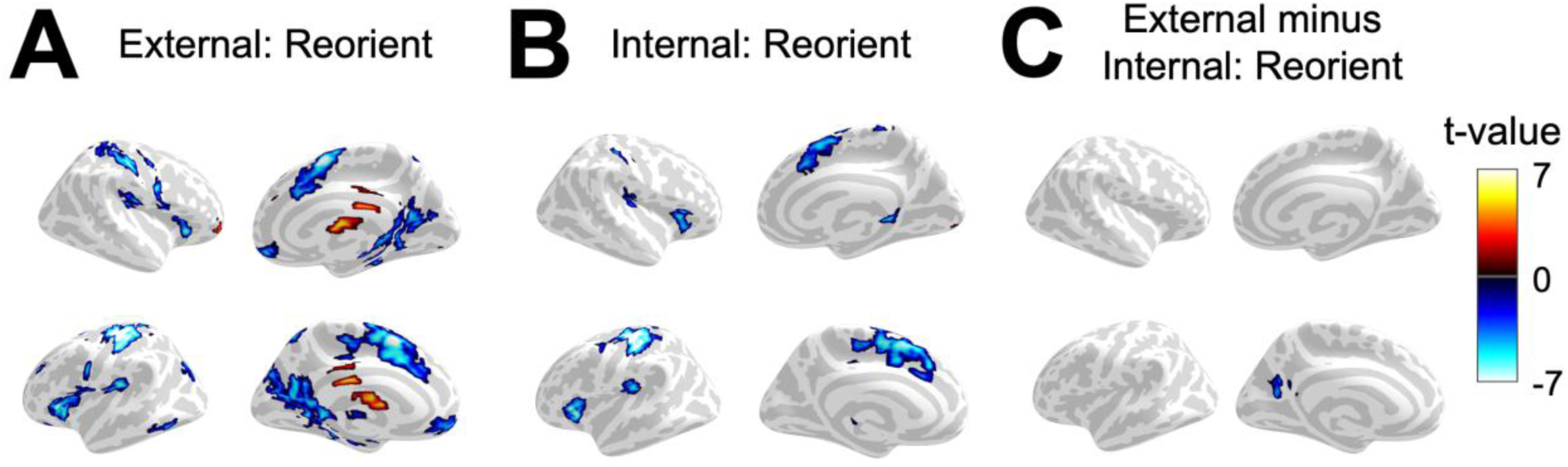
Voxel-wise univariate analysis for task periods involving attention reorienting. The group-level GLM estimated activation map (regression coefficients) for the 3 s period following the button press was extracted and projected onto the cortical surface. For panels A and B, positive t-values indicate voxels more strongly activated for the reorienting period and negative t-values indicate voxels that were less so; for panel C, positive values indicate voxels more active for the external compared to internal condition, while negative values indicate the opposite. **A.** The activation map for the meta-awareness period in the external condition. **B.** The activation map for the meta-awareness period in the internal condition. **C.** Subtracting the activation map for meta-awareness in the external condition from the period of meta-awareness in the internal condition.

## 4 Discussion

Our study asked whether human thought is naturally tuned towards a default internal state of cognition, called an internal bias of attention, or whether a similar automatic state can exist for external attention. We evoked both an external and internal attention state and compared the strength of each using self-reported mind-wandering events, whole-brain neural synchrony, and functional connectivity. We also investigated the cortical networks involved by analyzing moments of meta-awareness, when attention was directed at the current state of attention, and moments when attention was reoriented towards internal or external tasks.

### 4.1 Evoking Default Attention States

We found that in both external and internal attention conditions, the behavioral evidence of attention on the movies (based on reported mind-wandering events) and the neural evidence of attention on the movies (based on synchrony to the external stimulus) was highest on the first repetition relative to the last repetition, suggesting that the external stimulus was most easily represented and hardest to ignore when it was novel. We suggest that this attention on the novel, external stimulus represented a ‘default’ externally oriented state. In this interpretation, default, externally oriented states exist provided the right context. Our findings are in some ways comparable to the literature on external-internal multitasking paradigms in which information maintained in working memory conflicts with or facilitates an external task (Olivers et al., 2011; Soto et al., 2008). Those previous studies measured transient conflict between internal and external attention with low dimensional stimuli; we dynamically assessed internal-external intrusions over an extended period of time using more complex, ecologically valid stimuli.

We predicted that the strongest internal pull of attention would exist on the fourth repetition of the movie, producing a natural tendency to disengage from the external environment. This hypothesis was confirmed. According to the neural results, the external stimulus was least represented on the fourth repetition for both internal and external conditions, suggesting the least external attention. However, the results did include an interesting complexity. On the fourth repetition, there was still more mind-wandering when the task was to pay attention internally than when the task was to pay attention to the video. We had expected that four repetitions of attending to the same video would be more challenging than four repetitions of attending internally. Yet even after participants viewed the video four times, it was easier to pay attention to the movie than to ignore it and pay attention to one’s own breathing. Despite previous literature advocating for a prepotent internally oriented bias of attention (Verschooren & Egner, 2023), this finding shows a context in which an internal focus of attention is harder to maintain than an external one. These findings do not preclude that an internal bias exists in the broader context of day-to-day activities, but rather that there are some contexts, such as when engaged with movies (and perhaps also social media), when an external stimulus is exceptionally captivating, more so than internal simulation.

### 4.2 Mind-wandering

The term ‘mind-wandering’ has recently been criticized for its ambiguity, with some calling attention to the diverse range of definitions that vary by experimenter and task design (Christoff et al., 2018; Seli et al., 2018). Most commonly, mind-wandering has been defined as a shift in the contents of thought away from an ongoing task in the external environment to internally self-generated thoughts and feelings (Smallwood & Schooler, 2015). However, others have suggested that the definition of mind-wandering must involve a reference to intentionality (Dixon et al., 2014; Seli et al., 2016), as it can be influenced by volitional processes such as motivation (Seli et al., 2019) and can be intentional if the external task is not cognitively demanding (Robison et al., 2020).

The present study integrates both ideas, incorporating dominant external and internal attention states and their relationship to voluntary and involuntary cognition. Our findings are consistent with the view that mind-wandering can occur bidirectionally, from the internal to the external environment as well as from the external to the internal environment (Robison & Unsworth, 2015). Our findings also help to illustrate the link between mind-wandering, which involves spontaneous thought, and focused attention meditation, which involves controlled internal thought (Antoine Lutz et al., 2008). In our task, we operationalize mind-wandering as involuntary attention on items that are off task, regardless of the internal-external distinction.

Previous work manipulated the propensity toward mind-wandering by scaling task difficulty, finding that less mind-wandering events occur as the demands of the task increase and more mind-wandering events occur as the task demands decrease (Seli et al., 2016). In the present study, instead of experimentally modifying task difficulty, we modified the salience of the external stimulus. In previous literature, salience is often manipulated by changing low level features such as the contrast of neighboring pixels (Itti & Koch, 2001) or the overall informational value of a stimulus (Gottlieb, 2012). Here we used both the intrinsic interest value of the movie stimulus, and repetition of the stimulus, to alter its salience. Repetition is known to evoke anticipatory responses in learning paradigms (Michelmann et al., 2021; Richardson & Saxe, 2020; Turk-Browne et al., 2010) and weaker responses (i.e. stimulus suppression) when no explicit objective is specified (Aly et al., 2018; Grill-Spector et al., 2006). In our study, repetition effects occurred in the absence of a learning objective, for naturalistic, multi-modal stimuli.

We believe our study aligns with a previous suggestion that researchers should work to characterize the dynamics of mind-wandering rather than simply trying to define it (Christof et al., 2018). Although we operationalized mind-wandering in our experiment in a particular way, our purpose is not to champion one definition over another. Instead, we probed the dynamics of mind-wandering during internal or external focus, showing a graded scale along which mind wand wandering events increased or decreased. We found that prolonged internal focus induced more mind-wandering than prolonged external focus.

### 4.3 Controlled and automatic processing

It has been suggested that the distinction between automatic and controlled processing is suspect. Processing may vary continuously along an automatic-to-controlled scale, rather than in a discrete manner (Christof et al., 2018). We provide novel support for this idea. In our external attention condition, we found that automatic, external attention to the movie gradually waned with each repetition (based on the ISC analysis), and, in turn, the necessity for controlled processing increased (based on the connectivity analysis). Other studies have identified the relative contributions of individual and shared processing using these two fMRI analysis methods (Mochalski et al., 2024; Simony et al., 2016). Our study specifically investigated the time course of attention and attentional control mechanisms.

### 4.4 Meta-awareness

The relationship between meta-awareness and mind-wandering is primarily explored in mindfulness and meditation research. Meta-awareness is associated with increased control (Debettencourt et al., 2015; Jha et al., 2007), decreased mind-wandering events (Christoff et al., 2009; Levinson et al., 2014; MacLean et al., 2010; Mrazek et al., 2012), and increased ability to deflect negative internal thought (Deng et al., 2014; Nolen-Hoeksema et al., 2008; Teasdale et al., 2002). These studies have generally tested meta-awareness towards internal or external contents, without comparing the two. Studies that directly compare them focus on unimodal meditation objects that do not dynamically change (Stawarczyk et al., 2011; Weng et al., 2020).

The present study provides further evidence that meta awareness and control can alter internal and external modes of cognition in a dynamically changing environment (Kawashima et al., 2023; Schooler et al., 2011).

Networks involved in meta-awareness, and metacognition more broadly, are commonly reported in midline and orbital prefrontal structures (Christoff et al., 2009; Fleming & Dolan, 2012), but also bilaterally in the Insula (Christian et al., 2025; Hasenkamp et al., 2012). Here we found some activations outside of these canonical regions, including areas in the DMN and Control networks. These regions may play a more pronounced role in meta-awareness than previously thought. Alternatively, these activations may have been due to other cognitive (e.g., mind-wandering) and sensory-motor control processes occurring before the button press.

### 4.5 Neural signatures of Internal and External attention

Cortical networks are often characterized along an internal-external axis (Fox et al., 2015; Menon & D’Esposito, 2022). The DAN is associated with activation during goal directed external attention, and deactivation during internally oriented attention (Corbetta et al., 2008; Szczepanski et al., 2013). In contrast, the DMN deactivates when external attention is engaged and activates when internal attention is required (Fox et al., 2005; Shulman et al., 1997). Similarly, positive correlation between the FPCN and DAN is a signature of goal-directed external attention, and anticorrelation between the FPCN and the DMN a hallmark of internal attention (Dixon et al., 2018; Fox et al., 2005; Maillet et al., 2019; Spreng et al., 2010).

However, in the present study, the results did not support a simple internal-external distinction. When we analyzed moments of attention reorientation, attention networks were similarly engaged during shifts of attention toward external and toward internal content. When we analyzed attention control over time, networks traditionally associated with *external* sensory processing, including the DAN, remained significantly correlated with the FPCN in the *internal* attention condition, often more so than in the external condition. Similarly, the R TempPar, often associated with *internal* attention, was still significantly connected to the FPCN when the focus of attention was directed *externally*. This pattern indicates that networks traditionally associated with external, sensory processing or with internally oriented cognition may serve functional roles beyond the internal–external dichotomy, potentially reflecting more general processes such as sustained effort (Langner & Eickhoff, 2013; Sadaghiani & D’Esposito, 2015). Taken together, our work suggests that the distinction between internal-external attention networks may be less pronounced than previously assumed.

### 4.6 Limitations

We manipulated salience in the external condition by displaying movies that participants judged to be more or less exciting. In contrast, internal salience was held constant: participants were instructed to attend to their breath across all four repetitions, rather than to alternate among various internally focused tasks. Future works should aim to manipulate the internal task as well. For instance, instructing participants to focus on an autobiographical memory or simulating a future scenario – processes that naturally occur during unconstrained internal mentation (Baird et al., 2011; Kvavilashvili & Rummel, 2020) – may elicit more engagement with the internal environment compared to the external one. Enhancing internal salience in this way could weaken the dominance of the default externally oriented state evoked in our paradigm and help delineate the contexts in which internal and external modes of cognition compete (Dixon et al., 2014).

It is worth noting that mind-wandering during external attention is not always driven by spontaneous internal thought, nor is mind-wandering during internal attention necessarily caused by external distractions. When the task involves focusing externally, off-task episodes may arise not only from internal cognition (e.g., memories or thoughts) but also from competing external distractors in the environment (e.g., scanner noise). Conversely, when attention is directed inward, mind-wandering may stem from spontaneous internal mentation as well as from external sensory inputs. We did not have participants distinguish between mind-wandering sources as others have done (Robison & Unsworth, 2019; Stawarczyk et al., 2011).

We do not claim that our experiment, relying on participants’ own introspection and judgement, is the most accurate way to capture the phenomenon of mind-wandering. Our goal in employing the self-caught method, rather than alternative paradigms with stronger validation (e.g. the probe-caught method; Weinstein, 2018), was to elicit sustained meta-awareness and attentional control without interrupting the external stimulus. To this end, the self-caught method was suitable for the present design. However, other methods of measuring and manipulating mind-wandering may be useful in future designs.

## Author Contributions

IRC and MSG conceived the project, designed the method, and prepared the original draft. IRC collected the data and performed formal analysis. SAN advised on experimental design, methods, and reviewed and edited the original draft. LKK provided support with data visualization and analysis.

## Data Availability

All anonymized data are publicly available online (https://github.com/isaacrc/metaawareness-and-default-states)

## Acknowledgements

Supported by Princeton Neuroscience Institute Innovation Fund 24400E2349FA010 and by AE Studios grant 24400B1459FA010.

## Declaration of interests

The authors declare no competing interests.

## Notes

https://github.com/isaacrc/metaawareness-and-default-states

### Competing Interest Statement

The authors have declared no competing interest.

### Summary of Updates

This is a revised manuscript, including new results and discussion.

