## Supplementary material for "Meta-awareness, mind-wandering, and the control of ‘default’ external and internal orientations of attention": Supplment 1

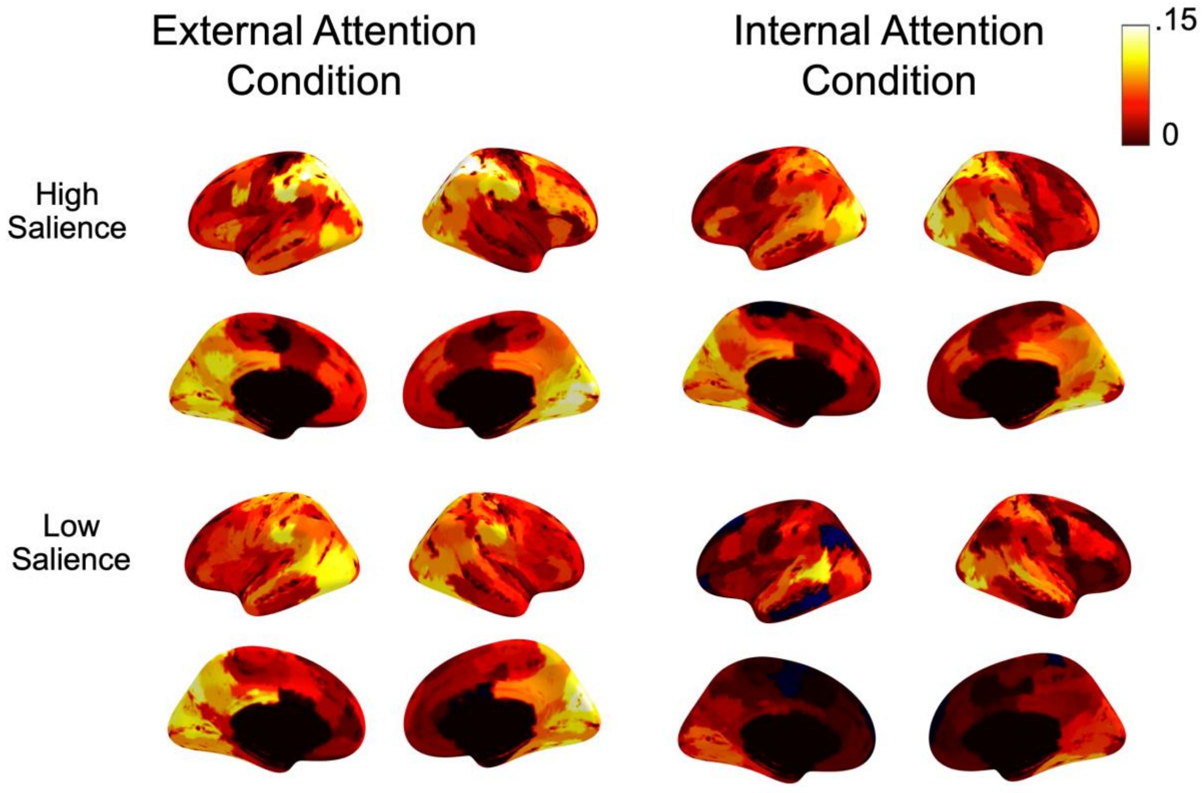


Figure 1. Difference in unthresholded mean ISC values when subtracting repetition 1 from repetition 4 across 200 brain regions. Larger decreases in ISC from repetition 1 to repetition 4 are illustrated in warmer colors. Darker regions show less change in ISC over time. The left column identifies differences when attention is directed externally and the right column shows when attention was directed internally. Difference values are averaged across high salience (top row) and low salience (bottom row) movies.

*Table 1*. Meta-awareness (0-3s before button press)

| Contrast | Harvard-Oxford Label | x | y | z | Peak Z |
|---|---|---|---|---|---|
| External-Internal | Paracingulate Gyrus | 3.5 | 16.27 | 34.3 | 5.51621588 |
| External-Internal | Orbital Frontal Cortex | -24.41 | 52.33 | -10.27 | 4.74349778 |
| External-Internal | Lateral Occipital Cortex, inferior division | 14.89 | -65.74 | 8.34 | -3.6232985 |
| External-Internal | Inferior Temporal Gyrus, posterior division | -47.37 | -50 | -17.17 | -3.6296438 |
| External-Internal | Lateral Occipital Cortex, superior division | -37.18 | -84.24 | -19.77 | -3.6561956 |

| | | | | | |
|---|---|---|---|---|---|
| External-Internal | Supracalcarine Cortex | -18.09 | -89.33 | -4.01 | -3.66054 |
| External | Paracingulate Gyrus | 8.77 | 14.76 | 37.2 | 14.444806 |
| External | Parahippocampal Gyrus, anterior division | 17.82 | -23.3 | -14.23 | 12.7566953 |
| External | Background | -49.42 | -54.71 | 64.14 | 10.4788943 |
| External | Background | -31.28 | 19.76 | 88.9 | 10.3978196 |
| External | Cuneal Cortex | -26.89 | 5.63 | -14.92 | 10.365106 |
| External | Angular Gyrus | -18.36 | -68.24 | 37.48 | 9.04989922 |
| External | Subcallosal Cortex | -10.53 | 48.25 | -3.63 | 8.93138156 |
| External | Background | -11.93 | -41.2 | 108.59 | 8.62684406 |
| External | Background | -32.1 | -18.26 | 32.27 | 8.6068535 |
| External | Middle Temporal Gyrus, posterior division | -57.15 | -55.27 | -6.39 | 7.72402386 |
| External | Background | -8.78 | 46.34 | 76.62 | 6.86725225 |
| External | Background | -2.41 | -47.6 | -34.99 | 6.85957754 |
| External | Background | -64.71 | 6.19 | 91.07 | 6.62402631 |
| External | Background | -14.62 | -17.42 | 109 | 5.74846858 |
| External | Cingulate Gyrus, anterior division | -7.59 | -25.57 | 34.81 | 5.6426425 |
| External | Superior Frontal Gyrus | -31.24 | 11.03 | 56.19 | 5.57809742 |
| External | Cingulate Gyrus, posterior division | 2.98 | -53.54 | 49.62 | 5.0987606 |
| External | Frontal Operculum Cortex | -31.56 | 8.75 | 17.57 | 5.05183802 |
| External | Temporal Fusiform Cortex, posterior division | -39.96 | -61.22 | -22.08 | -3.621811 |
| External | Background | -39.45 | -58.35 | 97.08 | -3.6224151 |
| External | Parahippocampal Gyrus, posterior division | 1.3 | -75.07 | 5.06 | -3.623861 |
| External | Background | -0.39 | -4.06 | 101.97 | -3.6262142 |
| External | Angular Gyrus | 15.24 | -60.27 | 71.12 | -3.628873 |
| External | Inferior Frontal Gyrus, pars opercularis | 18.09 | -24.32 | 43.53 | -3.6305018 |
| External | Background | -61.68 | -76.95 | -10.74 | -3.636219 |
| External | Background | -32.88 | -89.48 | 72.84 | -3.6378414 |
| External | Inferior Frontal Gyrus, pars opercularis | 13.41 | -22.19 | 79.11 | -3.6403602 |
| External | Insular Cortex | 17.82 | -10.97 | 69.8 | -3.642987 |
| External | Background | -22.1 | 39.14 | 94.98 | -3.6565902 |
| External | Inferior Frontal Gyrus, pars opercularis | 27.54 | -22.02 | 66.88 | -3.6689101 |
| External | Background | -18.21 | 14.45 | 17.99 | -3.7499133 |
| Internal | Subcallosal Cortex | 9.69 | 13.18 | 37.6 | 12.4357557 |
| Internal | Background | -3.98 | -10.42 | -14.04 | 11.9889356 |

| | | | | | |
|---|---|---|---|---|---|
| Internal | Background | -30.48 | 18.7 | 88.61 | 10.8785722 |
| Internal | Background | -30.06 | -18.9 | 32.38 | 10.4600478 |
| Internal | Subcallosal Cortex | -8.14 | 47.01 | -1.3 | 8.93872566 |
| Internal | Angular Gyrus | -28.5 | -67.7 | 46 | 8.67346053 |
| Internal | Background | -8.99 | 47.62 | 76.86 | 8.24367524 |
| Internal | Background | -4.36 | -45.58 | -35.32 | 7.63666718 |
| Internal | Middle Temporal Gyrus, posterior division | -55.69 | -54.67 | -9.39 | 7.16639351 |
| Internal | Out of bounds | -11.56 | -41.96 | 110.32 | 6.98613681 |
| Internal | Background | -64.1 | 4.18 | 89.9 | 5.96865509 |
| Internal | Cingulate Gyrus, anterior division | -7.38 | -23.78 | 37.32 | 5.89514817 |
| Internal | Background | 10.9 | 3.33 | 88.38 | 5.52170129 |
| Internal | Cingulate Gyrus, anterior division | 5.1 | -24.9 | 25.18 | 5.19020141 |
| Internal | Superior Parietal Lobule | -64.49 | -26.92 | 34.16 | 5.04579364 |
| Internal | Background | 9.22 | 17.19 | 4.72 | -3.6236263 |
| Internal | Parahippocampal Gyrus, posterior division | 0.91 | -76.49 | -10.2 | -3.6247497 |
| Internal | Background | -40.29 | 50.22 | 37.25 | -3.6364961 |
| Internal | Out of bounds | -47.67 | -7.94 | 112.43 | -3.6415124 |
| Internal | Background | -7.52 | -64.38 | 97.2 | -3.6573148 |

*Table 2.* Reorienting (0-3s after button press)

| Contrast | Harvard-Oxford Label | x | y | z | Peak Z |
|---|---|---|---|---|---|
| External-Internal | Cingulate Gyrus, posterior division | -14.91 | -59.22 | 21.53 | -3.6283203 |
| External | Insular Cortex | 19.76 | 34.23 | 40.06 | 8.52121842 |
| External | Background | -17.44 | -3.4 | 27.89 | 7.7371325 |
| External | Background | -50.03 | -81.62 | 69.35 | 7.2578854 |
| External | Supracalcarine Cortex | -1.74 | -90.95 | 39.57 | 6.64754054 |
| External | Background | -12.19 | 40.49 | 38.5 | 6.35997605 |
| External | Cingulate Gyrus, anterior division | -6.5 | -24.93 | 37.21 | 6.21762746 |
| External | Background | -5.36 | 59.7 | 63.41 | 6.11157213 |
| External | Background | -31.74 | 53.25 | 90.21 | 5.97870669 |
| External | Background | -32.93 | -101.37 | 38.71 | 5.71929701 |
| External | Lateral Occipital Cortex, superior division | -53.49 | -79.08 | -3.63 | 5.51898563 |
| External | Parahippocampal Gyrus, anterior division | 18.05 | -25.16 | -10.15 | -3.6218116 |
| External | Planum Polare | -47.4 | -16.09 | 6.65 | -3.6218602 |
| External | Paracingulate Gyrus | 12.08 | 4.41 | 34.06 | -3.622011 |

| | | | | | |
|---|---|---|---|---|---|
| External | Background | 18.93 | -26.74 | 87 | -3.6220902 |
| External | Lateral Occipital Cortex, inferior division | 8.89 | -72.66 | 7.68 | -3.6224511 |
| External | Temporal Fusiform Cortex, posterior division | -42.24 | -56.47 | -28.12 | -3.6226599 |
| External | Out of bounds | -14.02 | 3.35 | 110.57 | -3.6227391 |
| External | Postcentral Gyrus | -22.54 | -55.37 | 35.74 | -3.6235547 |
| External | Background | -14.62 | -22.92 | -14.41 | -3.6265359 |
| External | Background | -7.7 | 4.2 | -36.18 | -3.6271112 |
| External | Angular Gyrus | 21.1 | -65.1 | 47.02 | -3.6273443 |
| External | Frontal Pole | -28.3 | 19.4 | -7.58 | -3.6392687 |
| External | Background | -28.8 | -21.28 | 25.09 | -3.6413343 |
| External | Background | -39.46 | 47.86 | 34.19 | -3.6425497 |
| External | Background | -29.55 | 26.65 | 74.5 | -3.6472394 |
| External | Background | -14 | -21.88 | 92.66 | -3.6476029 |
| External | Heschl's Gyrus (includes H1 and H2) | -45.34 | -36.16 | 6.68 | -3.6800152 |
| External | Paracingulate Gyrus | -2.99 | 37.57 | -2.2 | -3.6934057 |
| Internal | Insular Cortex | -16.16 | -1.59 | 56.5 | 6.04041175 |
| Internal | Background | -14.74 | -8.16 | 18.61 | 5.76841163 |
| Internal | Inferior Frontal Gyrus, pars opercularis | -32.23 | -9.19 | 72.27 | 5.45952105 |
| Internal | Background | -6.29 | 63.54 | 58.88 | 4.98045625 |
| Internal | Lateral Occipital Cortex, superior division | -52.46 | -80.48 | -13.01 | 4.93219535 |
| Internal | Background | -45.64 | -92.32 | 59.53 | 4.88009019 |
| Internal | Background | 14.16 | 44.54 | 46.24 | 4.70680567 |
| Internal | Paracingulate Gyrus | 12.05 | 5.55 | 32.56 | -3.6220326 |
| Internal | Background | 17.36 | -24.66 | -10.93 | -3.6253476 |
| Internal | Background | -14.42 | -23.43 | -16 | -3.6284277 |
| Internal | Inferior Frontal Gyrus, pars opercularis | -14.49 | -23.19 | 85.92 | -3.6299481 |
| Internal | Insular Cortex | 25.6 | 1.99 | 53.91 | -3.6304978 |
| Internal | Inferior Temporal Gyrus, temporooccipital part | 25.51 | -34.95 | 49.59 | -3.635742 |
| Internal | Frontal Pole | -28.06 | 23.97 | -2.92 | -3.6358121 |
| Internal | Background | 9.06 | -25.48 | 93.18 | -3.6387819 |
| Internal | Frontal Pole | -29.08 | -24.42 | 20.4 | -3.6528618 |
| Internal | Supramarginal Gyrus, anterior division | -48.55 | -50 | 57.05 | -3.6621735 |
| Internal | Background | -27.65 | 25.36 | 76.79 | -3.6733896 |

| | | | | | |
|---|---|---|---|---|---|
| Internal | Inferior Temporal Gyrus, temporooccipital part | -27.04 | -38.04 | 54.02 | -3.7048262 |
